# Effect and Mechanism Analysis of Melittin on Promoting Muscle Differentiation of Adipose Stem Cells

**DOI:** 10.1101/2025.06.05.658180

**Authors:** Ming-Jun Li, Yu-Xia Yang, Jian Zhang, Xiang-Ji Meng, Wen-Kang Liu, Shuo-Guo Wang, Dian-Wei Liu, Meng-Bo Dang, Jian-Yang, Xiao-Mei Dai, Tao-Bao, Ya-Wang, Wen-Yong Fei

## Abstract

**Purpose:** Melittin is the main peptide component of melittin venom, with anti-allergic, anti-inflammatory, anti-arthritis, anti-cancer, and neuroprotective effects. Melittin has been confirmed to have anti-inflammatory effects in immune and liver cells and is widely used in treating arthritis and rheumatic diseases. At the same time, related studies have shown that melittin can enhance the expression of muscle differentiation related factors in mice. However, whether the melittin can improve the expression of stem cells into muscle differentiation and use it to treat Skeletal muscle atrophy, it has not yet been clarified.As "seed cells," stem cells have been widely concerned in the field of regenerative medicine research, such as embryonic stem cells, induced pluripotent stem cells, mesenchymal stem cells, etc., among which adipose mesenchymal stem cells (ADSCs) have excellent application potential in the treatment of Skeletal muscle atrophy because of their quantitative advantages, easy access, and inherent good differentiation potential and self-renewal ability. To investigate whether melittin can stimulate ADSCs MYOGenic differentiation and whether combined with ADSCs can treat Skeletal muscle atrophy.

**Methods:**

1. Four concentration ranges of melittin were set, namely Group A (20 mg/L, 40 mg/L, 80 mg/L, 160 mg/L, 320 mg/L), Group B (2 mg/L, 4 mg/L, 8 mg/L, 16 mg/L, 32 mg/L), Group C (0.2 mg/L, 0.4 mg/L, 0.8 mg/L, 1.6 mg/L, 3.2 mg/L), and Group D (0.02 mg/L, 0.04 mg/L, 0.08 mg/L, 0.16 mg/L, 0.32 mg/L), Intervention of adipose stem cells with different concentrations of melittin for 48 hours, followed by measurement of relative cell viability using the CCK-8 method.
2. Melittin after 48 hours of intervention with ADSCs, Western blot was used to detect the expression levels of MYOGenic differentiation related factors.
3. The control group (C), melittin group (M), dexamethasone group (D), and Dexamethasone+Melittin group (D+M) were set separately. After 12 hours of intervention with dexamethasone (10umol/L), the ADSCs were added with melittin for 48 hours. Western blot was used to detect MYOGenic differentiation related factors MYOGenic(MYOG), MYF5, MYOD, MYH4 p38 and p-p38 protein. In order to observe the process of melittin repairing cell atrophy, detected MYOD and MYOG using immunofluorescence.
4. Randomly divided into control group (C), damage group (T), stem cell therapy group (SC), melittin combined with ADSCs therapy group (SC+M). Group C does not do any treatment. Group T, SC, and SC+M set up a mouse shoulder sleeve injury model, all from the left gang of the rats, and the suture is marked with surgical sutures. On the first day of surgery, The SC group of the muscle injecting ADSCs(1×10^6^)in the SC group, and the ADSCs after the melittin intervention of the SC+M mouse is repeatedly treated every seven days. On the left, the tendon is extracted, and the muscles are extracted to perform related tissue-related testing such as Su Mu Jing-Yi Hong dyed (H&E dyed), immunohistochemistry, muscle fiber weight and diameter.

**Results:**

1. CCK-8 Experiment shows that the suitable concentration of Melittin has no significant toxic effect on ADSCs.
2. Western blot experiment shows that the appropriate concentration of Melittin can enhance the expression of muscle differentiation related factors in ADSCs.
3. Western blot and immunofluorescence staining results show that the expression related factors of the D+M group are better than that of group D, and the expression related factors of forming muscle differentiation in group M is also better than group C.
4. Melittin can promote ADSCs muscle differentiation through the p38-MAPK signaling pathway.
5. Animal experiments have shown that the expression related factors of the SC+M treatment group’s muscle differentiation related factors is significantly higher than that of the SC therapy group, and the expression related factors of the two groups of rats of SC+M and SC groups are better than the T group Rat. Organic research has confirmed that rats who have been treated with ADSCs treated with melittin have significantly improved in muscle weight, muscle fiber diameter, and muscle fiber cross-sectional area. At the same time, the result of immunohistochemicals shows that the expression of MYOD and MYOG in the SC+M group is better than the SC group.

**Conclusion:** The results of this study indicate that melittin can promote MYOGenic differentiation of ADSCs and increase the expression of MYOGenic differentiation related factors in atrophic ADSCs.Secondly, melittin can activate the p38-MAPK signaling pathway and promote the MYOGenic differentiation of ADSCs. Stem cells treated with melittin intervention can improve the recovery of muscle mass, muscle fiber diameter, and muscle fiber cross-sectional area in rats with skeletal muscle injury compared to rats treated with stem cells alone. Therefore, the above results provide insights into MYOGenic differentiation and may provide a potential alternative strategy for related diseases such as skeletal muscle atrophy.

## Introduction

Skeletal muscle is one of the largest, most dynamic and plastic tissues in the body, accounting for 40%of the total quality of the body, which is crucial to exercise and energy metabolism ^[1]^. The loss of skeletal muscle quality and strength, such pathological phenomena is called skeletal muscle atrophy ^[2]^. Skeletal muscle atrophy damage the body’s resistance to stress and chronic diseases, seriously reduce people’s quality of life, increase the incidence and mortality ^[3]^. At present, rotating cuff damage is the most common in the field of sports medicine. Although the clinical effect of surgery is satisfactory, there is still a problem with the recovery of the shoulder sleeves. One of the main reasons for skeletal muscle atrophy of the main reason is that the continuous muscle weakness is weak ^[4]^. Although there are many methods for treating skeletal muscle atrophy, it still cannot fundamentally relieve and control the progress of the disease ^[5-6]^. Therefore, the treatment of skeletal muscle atrophy is constantly exploring the treatment of skeletal muscle atrophy, increasing clinical treatment level, and a long way to go.

With the development of stem cells, the importance of stem cells to skeletal muscle disease is discovered. Stem cells are widely concerned as "seed cells" in the field of regeneration medicine. For example, embryonic stem cells, inducing polyo-stem cells, and germinated stem cells ^[7]^. Among them, lipid skeletal muscle atrophy shows excellent application potential ^[8]^. However, related studies have shown that only 15%of stem cells harvested from extracted from extracts can be fully differentiated. At present, researchers are trying to promote stem cells into muscle division, such as drug induction, physical induction, cell stent induction, Wait, although various methods show that it can promote ADSCs muscle differentiation, there are still problems such as low induction efficiency, the induction method significantly inhibiting the self-renewal ability of stem cells, and low survival rate after stem cell transplantation ^[7]^.

Curcumin, Korean medicine and other traditional Chinese medicine ingredients have achieved good results in treating skeletal muscle atrophy. The mango flower yellowin can inhibit skeleton’s atrophy by promoting cellular derivative differentiation. Therefore, traditional Chinese medicine has become increasingly important in the treatment of skeletal muscle atrophy, and has become a research hotspot^[9-11]^. Melittin is the most prominent ingredient in bee toxin. It is a two-family hexagon peptide, which accounts for about half of the dryness of bee toxin and belongs to a traditional Chinese medicine ^[12]^. Studies have found that melittin have significant effects in the fields of anti-inflammatory, antioxidant, analgesic, sterilization, anti-tumor and anti-radiation ^[13]^, and used to treat diseases such as skin and multiple sclerosis ^[14-15]^. Related studies have shown that melittin can improve the expression of muscle differentiation related factors in mice skeletal muscle cells [16], but whether it can act on ADSCs and improve ADSCs muscle differentiation efficiency, it still needs to be verified.

In this study, it can explore whether the melittin can promote ADSCs muscle differentiation through cell experiments, and at the same time establish a cell atrophy model to explore whether the melittin can increase the expression of the muscle differentiation related factors in ADSCs. And establish a mouse skeletal muscle atrophy model, evaluate the treatment effect of ADSCs after melittin intervention in treating skeletal muscle atrophy of rats, trying to find a way to enhance ADSCs activity, promote ADSCs muscle division, improve ADSCs treatment of skeletal atrophy of skeletal muscle atrophy A new method.

## Materials

### 1.1 Experimental animal

Buy male SD rats (6-8 weeks, 240 ±10g, 12 in each group) from Wukong Biotechnology Co. Ltd. (China), and allows adaptation 7 days before the start of the study. SD rats are raised under standard conditions (12:12 hours of light dark circulation, 23 ±1 ℃), and free water and food. The University Animal Nursing and Use Committee approved all animal experimental plansat the Ethical Committee of the SuBei People’s Hospital (acceptance number: 2021ky-001). All experimental procedures using animals are based on the "Ethical Review Methods involved in human biomedical research", "Regulations on the Management of Quality Management of Drug Clinical Test, "Regulations for the Management of Medical Device Clinical Test Quality Management", international ICH-GCP and other laws and regulations and ethical specifications conduct.

### 1.2 Reagent

Melittin (Shanghai Yuanye Biotechnology Co. Ltd.), Demomeson (Beijing Solibao Biotechnology Co. Ltd.), PBS (Beijing Solabao Biotechnology Co.Ltd.), DMEM medium (US GIBCO Company), fetal cow Serum FBS (American GIBCO Company), penicillin-chaincin (Shanghai Biyuntian Biotechnology Co.Ltd.), Malaysia (American GIBCO Company), CCK-8 cell vitality detection kit (American Simer Flying Shier Technology Company), MYOD antibodies (American Simer Flying Shier Technology Co.Ltd.), MYOG antibody (American Simer Flying Shier Technology Co.Ltd.), MyHC antibody (American Simerfield Technology Company), MYH4 antibody (American Simerfield Technology Company), IGG (Beijing Bolossen Biotechnology Company), BCA protein testing kit (Jiangsu Kiki Biotechnology Co.Ltd.)

### 1.3 Experimental Equipment

Super Pacific Workshop (Suzhou Purification Equipment Factory), carbon dioxide training box (German HERAEUS), high-speed centrifugal machines (German Leica), micro-centrifugal machines (Shanghai Precision Experimental Equipment Co., Ltd.), Analysis Library (Germany Saidis Group), High-pressure disinfection cooker (Shanghai Shanghai Honorary Trading Co., Ltd.), refrigerator (China Hisense Company), constant temperature water bath (German Leica), Guli (Wuhan Seville Biotechnology Co., Ltd.) Er Technology Co., Ltd.), enzyme standard (Saimo Flying Shier Technology Company), transmitted electronics microscope (Japanese Hitachi), ordinary optical microscope (Nikon in Japan), ordinary optical microscope (Suzhou Sienc Industrial Technology Co., Ltd.), Fluorescent microscope (Japan Olympus Optical Technology Company), X-ray diffraction meter (Japanese Rigaku), co-focused laser scanning microscope (German Leica), electrophoretes (Hercules Technology Company), gel imaging system (USA Hercules Technology Company).

## Research methods

### 2.1 separation and extraction of ADSCs

All experiments were approved by the Animal Ethics Committee (acceptance number 2021-ky001) and conducted in accordance with regulations and guidelines. SCS’s preparation is based on the method used by predecessors. Anesthesia SD white rats under sterile conditions, cut 3cm in parallel at the groin of SD rats, separate the SD rats, the subcutaneous fat tissue of the groin, and rinse the SD rats with DPBS twice with DPBS. The vascular and connective tissues are eliminated with micro and the adipose tissue is cut into small blocks with ophthalmology, and the size is about 1 mm³. The cut SD rats are used for digestion of 0.2%IA collagen enzymes of an equal volume, and then the water bath is digested 1.5h (37℃) to shake the digestive adipose tissue intermittently to fully digest it. Then use the equivalent volume to completely medium (90%DMEM-F12+10%fetal beef serum+concentration of 1%double resistance) to terminate the digestion of pancreatial enzymes, 200 mesh screen filtration, 1000 R/min centrifuged 5 min, then suck out the Qingqing Qingqing Liquid, continue to add DPBS heavy suspension cells 1000 R/min centrifuged 5 min. Use a complete cultivation solution to the suspension cells, inoculate in the 25 mm² culture bottle, place ADSCs at 37℃, and cultivate 48 h in the 5%CO_2_ incubation box. Change the medium once every 2 days, and observe the form of ADSCs under the inverted microscope.

### 2.2 The cultivation and transmission of ADSCs

Treat the cells in the growth medium and in the cell culture box, and the interval between 2 to 3 days is performed. Pay attention to observing the cell morphology and cell density of ADSCs. When the density reaches 70%to 80%, it is passed on. Use the liquid in the absorbent dine before the cell transmission, wash it with PBS solution, and then use 0.25%pancreatase digestive solution to digest cell digestion. After the cells fall off, 1000r/min is centrifuged for 5min. : The ratio of 3 ∼ 1: 4 is passed on, and the marks are made in the incubation box for training.

### 2.3 The freezing and recovery of ADSCs

Prepare the cell gradient frozen box in advance, configure the frozen storage liquid in the ratio of 1: 9 and place it in a 4℃ refrigerator to pre-cold. After the ADSCs cell sample is prepared to be perfect, use 0.25%pancreatase digestion cells, suck ADSCs into a 15ml centrifugal tube centrifugal tube, and suck the upper liquid in the centrifugal tube. Add PBS heavy suspension cells. Put the cell gradient frozen storage box, immediately store it in a low-temperature refrigerator at -80℃, and transfer it to the liquid nitrogen frost the next day. When a cell recovery is needed, preheat the water bath in advance to 37℃, remove the frozen storage tube from the liquid nitrogen, add the water bath and thaw, then disinfect the tube with alcohol, and move the cell suspension in the centrifugal tube to 15ml Sterile centrifugal tube, add the medium of foul cells, suck out the cells on the cell, re-use the medium suspension cells, and then transfer it to the new medium and put it in the incubation box to cultivate the medium the next day.

### 2.4 ADSCs form identification

Take the stable transmission, the growth density of more than 50 % and the good state of ADSCs, uses an optical microscope (CKX53, Olympus, JAPAN) to observe the ADSCs pattern and takes the records.

### 2.5 Analysis of ADSCs cell activity and best concentration analysis

Take 4 bee toxic peptide concentration interval, group A (20 mg/L, 40 mg/L, 80 mg/L, 160 mg/L, 320 mg/L), group B (2 mg/L, 4 mg/L, 8 mg/L, 16 mg/L, 32 mg/L), group C (0.2 mg/L, 0.4 mg/L, 0.8 mg/L, 1.6 mg/L, 3.2 mg/L), group D (0.02 mg/// L, 0.04 mg/L, 0.08 mg/L, 0.16 mg/L, 0.32 mg/L), melittin action ADSCs 48 hours, and then related testing.

Take the stable transmission, the growth density of more than 80 % and the good state of ADSCs, abandon the upper clearing, wash it with PBS 3 times, digest and collect cells in pancreat enzymes, re-suspension, count. Adjust the cell suspension, and finally make the number of cells per poron of 96 pores of about 5,000. Then put the 96-hole board in the culture box and incubate 24h to make the cells stick wall. Treat the cells according to Table 1, and set a blank group (cell-free), each group of 6 compound holes. After the processing is completed, abandon the 96-hole board in the middle of the Plains medium, and add 1.0×10^-4^L of the medium containing CCK-8 (CCK-8: Curma = 1: 10) per hole per hole. In the box, after 4h in the environment of 37℃, the OD value is measured using the enzyme marker (450nm using the light source wavelength). Collect data and calculate the proliferation vitality of each group of cells according to the following formula: cell proliferation vitality = [(Experimental OD value-blank group OD value)/(normal group OD value-blank group OD value)× 100 %].

### 2.6 Western-BLOT The effect of different concentration of bee toxin peptides on ADSCs muscle differentiation related factors

For cell samples, all cells are prepared on the seventh day; first extract protein, cell samples use centrifugal collection of cell precipitation, wash with PBS after the sample collection, and then use the pre-cold RIPA cracking solution to be split on the ice. The centrifugal tube is placed in a low-temperature and super-centrifugal machine for 12000r/min centrifuged for 10 minutes. The liquid clearance is transferred to the new EP tube, and then the BCA protein testing kit can be quantified by the BCA protein detection kit, and then the protein imprint is analyzed with the liquid clearance. The sample is stored in 20℃ for further use. Use 10% polyacrylamide gel with sodium sodium sodium sodium sulfate polyacrylamide gel electrophoretic (40 mg), and then use the semi-dry transfer system to the PVDF film. The printed traces are closed for 1 hour in the Tris buffer salt solution (TBS) containing a 0.1% Tween-20 Tris buffer (TBS). Subsequently, the membrane and the antibodies of the membrane with MYOG, MYH4, MYF5, and Gapdh protein were cultivated overnight at 4 °C. Finally, the anti-rabbit or anti-mouse IgG two-anti-bred anti-rabbit or mice IgG two-hour anti-mouse IgG two-hour anti-mice and mice are then washed for 10 minutes with TBS-T. Test. After testing, wash the membrane four times with TBS-T, wash each time for 5 minutes each time, and use Biotechnology (SHANGHAI, China, China, China) at 25℃ for 15 minutes to detect other proteins. Then use TBS-T to wash the film 4 times, 5% (W/V) to seal milk with 5% (w/v) each time, and perform immunohism.

### 2.7 Establishment of ADSCs cell atrophy model and cell experimental group

Take the stable transmission, the growth density of more than 80 % and the good state of ADSCs, at the same time add 10umol/L semimetisone to the 100ml medium to dilute, add the diluted dexamethasone to the ADSCs dine, induce ADSCs to shrink 12h, establish Cell atrophy model.

In order to explore the effects of melittin on the atrophic cells, the cell atrophy model is established through the dexamethasone atrophy, and the control group (C), the bee toxic peptide group (M), the dexamethasa group (D), the ground plug, the ground stuff, After intervention of ADSCs in the ground, the beexin peptide intervention is interfered with 48h, and the ADSCs is interfered with 48h after intervention of the ADSCs.

### 2.8 Western-BLOT detection of the expression of the correlation related factors in each group of ADSCs cell atrophy model

The above Western-BLOT detection method is used to detect the effects of MYOG, MYOD, MYH4, MYF5, Gapdh, and explore the effects of melittin on the expression of ADSCs in the group of ADSCs in each group of cell atrophy models.

### 2.9 The effect of immunofluorescence dyeing appraisal bee toxin peptide on the shrinking ADSCs muscle differentiation related factors expression

Place the cells on the ceiling of the lid. When the cell growth is fused to 95%-100%, it is placed in 37℃, and the 5%CO_2_ incubation box is cultivated. Wash 1×PBS 3 times, clean for 10 minutes each time, and then place the cleaned cells in 4%of the formaldehyde temperature for 20-30 minutes. Use 1 ×PBS 3 times again, 10 minutes each, and 0.2 with 0.2 each time. %Triton X-100 transparent for 2-5 minutes, washed 1×PBS 3 times, 10 minutes each time, 5%BSA room temperature was closed for 30 minutes, and one resistance (diluted with 1%BSA) was placed in a wet box, 4 degrees 4 degrees, 4 degrees 4 degrees After overnight, wash 1 ×PBS 3 times, 10 minutes each, add two-resistance (diluted with 1%BSA) for 30 minutes, close the light, wash 3 times 1×PBS, 10 minutes each time, 95%glycerin seal. The Zeiss LSM800 focuses on laser scanning microscope to shoot fluorescent images.

### 2.10 Western-BLOT The effect of bee toxin peptide on the p38-Mapk signaling pathway

The above Western-BLOT detection method is used to detect p38, p-p38, and Gapdh protein to explore whether the melittin can promote ADSCs muscle division through the p38-Mapk pathway.

### 2.11 Skeleton atrophy of the rats model establishment and animal experimental group

All rats are approved by the Animal Ethics Committee. Adopting conventional methods to divide 48 male SD rats into four groups: control group (C); Gangshandochylastation group (T); stem cell therapy group (SC); To. SD white rats are anesthesia with 2% iso fluoride and oxygen. After the skin is cut, cut it vertically on the bougainvillea, exposing the shoulder sleeve tendon at the shoulder joint. The tendon cutting group completely cut the upper lumpy tendon and lower gang tendon, removed some tendons, and sutured to reduce the chance of accidental healing. In the control group, the rats of rats were kept complete. The rats of the Gangshang muscle were from the Gangshamus muscle of the left shoulder sleeves from the left, but no treatment was made. The stem cell therapy group and bee toxin peptide combined with stem cell therapy group are separated from the left rotator sleeve Gangsomas. On the first day of the Okura muscle, ADSCs will perform muscle injection of the rats of the stem cell therapy group (about 1 ×106). At the same time The ADSCs muscle after intervention of bee toxin peptide intervenes the muscle injection (about 1 ×106) of the rats of the bee toxin peptide combined with the therapy group of the bee toxin peptide, and then inject ADSCs to the rats of the stem cell group every 7 days. The treatment of stem cells after the treatment of bee toxin peptide was established, and the chronic damage model of the rotator sleeves was established for 8 weeks. The mouse was held in the last week to collect and muscle samples.

### 2.12 Organization of tissue dyeing observation

1. Fixed and decapatically: After the separation of Mouse Gang’s muscle anatomy is separated, it is fixed for 24h in the 4%polymerization solution that is configured in advance. For amine four-ethyl acid (EDTA) solution for 2 days, the specimen is obtained for preliminary slices every 5mm along the cross section.
2. Desequent and transparent: After decontamination, place the good muscle specimens of the shareholders at 70%, 80%, 90%, and 100%concentration ethanol solutions, dehydrated for 30min in turn, and then placed in 50%diliacne, 75%2 two two two, 75%two two two. Toluene and 100%toluene are transparent.
3. Waxing and embedded: Preheat the thermostat box in advance, soak the transparent specimens with a mixture and pure paraffin solution of paraffin and dyshalne in turn, and pour the molten wax into the wax box. Put the specimen in the wax box and place it in iceCool it for 30 minutes, take it out after solidification.
4. Slice and grilled slices: Cut the wax block into small pieces with a blade, slices at 5mm thickness, place it at 45 °C warm water and wait for wax tablets to spread in the water., Bake slices in the oven to dry,Save at low temperature.

## Su Mu Jing-Yigan staining (H & E staining)

The produced slices are dehydrated in dysfoline, then placed at 100%, 90%, 80%, and 70%concentration ethanol solutions soaked 5min in turn, and finally rinsed with PBS for 5min. Drip the slices and add Suwu essence chromatography for 5min, rinse with distilled water, add a differential liquid 10s, rinse the water again, and add the blue fluid. After returning to the blue, put the slices into the red dyeing liquid dyeing for 3min and rinse it. Put slices in 70%, 80%, 90%, and 100%alcohol dehydration again, use dystershopine for transparent, seal with neutral resin after processing, cover the cover glass microscope (Nikon, Yangzhou, China) Observe the damaged muscle damage area on the dyeing glass under 400 times large mirror, and analyze the image through a microscope, and use ImageJ software to measure the muscle fiber cross sectional area CSA (total area of muscle fiber cross section/number of muscle fibers)^[17]^.

## Immunohistochemistry

Place the selected skeletal muscle tissue wax block on the cold platform. When the slices can be performed, the slicing machine is roughly sliced to make the tissue as complete as possible, then cut into a thin piece of 3 μm, placed in a 50℃ water bath pot Among the slices, after the slices are fully expanded, remove the special anti -off-carrier sheet for immunohistochemistry, so that the tissue is spread out and placed in a 70℃ oven for 40 minutes to melt the paraffin on the slices. Show dehydration: Take dehydration on slices, place it in the configuration-dehydrated dewaxing solution for 10min, and then place the gradient ethanol (concentration of 100%, 100%, 95%, and 75%, respectively. 2min, remove it in distilled water. Antigen repair: You need to determine whether and what method is taken according to the detection index; when using high-pressure repair, 1min30s is placed in the repair solution of the high-pressure environment. It is based on the PD-L1 immunohistochemical meal repair liquid to repair the antigen tissue. Dye: add the first one-resistant working liquid, incubate for 1h 37℃, rinse with PBS buffer 2-3 times after incubation; add the corresponding two-resistant working liquid, incubate for 25 minutes at 37℃, rinse with PBS buffer after incubation after incubation. 2-3 times; add color rendering, incubate for 3-5min, and rinse with PBS buffer 2-3 times after incubation. Repeated dye: Place 1min30s in the Sumuin solution and remove it, and rinse it with tap water for 2-3 times. Differential: Place 10s in the differential liquid, 1min30s in water; Blue: Place 1min30s in the anti-blue solution. (7) Dehydration: dehydrate in gradient ethanol and grill in the oven. (8) Seal: Drop a drop of neutral resin and sample slices, cover it with a glass slice. (9) Mirror inspection; observe under microscope to evaluate the dyeing situation of the slice tissue.

### 2.13 Rat skeletal muscle weight and fiber diameter assessment

After eight weeks, all rats were executed, separated on the left gangshando tendon, and used 0.01 to accurately electronic scale to say that the weight of Chonggang’s upper muscle and ruler (MM) measured the diameter of the muscle fiber.

### 2.14 Western-BLOT detection of related factors in animal tissue in animal Gangsomas

Rat Okamoto’s upper muscle was collected on the 8th week. The Okamoto tissue separates the muscles with surgical scissors and preserves the tissue at -80℃ until it is used. The frozen gangto tendon tissue crushed in the cooling mortar, and was uniform in the protein extracting solution (BioteChnology, SHANGHAI, China, CHINA), and cracked for 60 minutes on the ice. By rewarding cracks and recycling at 4℃ for 10 minutes at 13,000 RPM centrifuge. Then use the BCA protein testing kit to quantify the protein, and then analyze the protein marks with the liquid clearance. The sample is stored in 20 °C for further use. Use 10% polyacrylamide gel with sodium sodium sodium sodium sulfate polyacrylamide gel electrophoretic (40 mg), and then use the semi-dry transfer system to the PVDF film. The printed traces are closed for 1 hour in the Tris buffer salt solution (TBS) containing a 0.1% Tween-20 Tris buffer (TBS). Subsequently, the membrane and the antibodies of the membrane with MYOG, MYOD, MYH4, MYF5, and Gapdh protein were cultivated overnight at 4℃. Finally, the anti-rabbit or anti-mouse IgG two-anti-bred anti-rabbit or mice IgG two-hour anti-mouse IgG two-hour anti-mice and mice are then washed for 10 minutes with TBS-T. Test. After testing, wash the membrane four times with TBS-T, wash each time for 5 minutes each time, and use Biotechnology (SHANGHAI, China, China, China) at 25℃ for 15 minutes to detect other proteins. Then use TBS-T to wash the film 4 times, 5% (W/V) to seal milk with 5% (w/v) each time, and perform immunohism.

### 2.15 Statistical Analysis

All data is represented by the average ±standard deviation, and the comparison between the two groups through the double t -T-test analysis. The two-factor variance analysis (anova) and the DUNCAN multi-test are compared to multiple sets of data. P <0.05 is defined as a significant difference compared with the control group and is marked with*, P <0.01 is marked with **, P <0.001 is marked with ***, P <0.0001 **** mark. All data in this study is analyzed using Graphpad Prism 8.

## Results

### 3.1 Appropriate concentration of melittin has no significant effect on the activity of ADSCs

To explore the appropriate concentration of ADSCs for bee toxin peptide, set 4 bee toxin peptide concentration intervals, namely group A (20 mg/L, 40 mg/L, 80 mg/L, 160 mg/L, 320 mg/L),), Group B (2 mg/L, 4 mg/L, 8 mg/L, 16 mg/L, 32 mg/L), group C (0.2 mg/L, 0.4 mg/L, 0.8 mg/L, 1.6 mg/l, 1.6 mg/l, 1.6 mg/l, 1.6 mg/l, 3.2 mg/L), group D (0.02 mg/L, 0.04 mg/L, 0.08 mg/L, 0.16 mg/L, 0.32 mg/L), intervene After the CCK-8 method is used to determine the relative activity of cells. The experimental results show that under the intervention of ADSCs in the concentration intervention of the A bee toxin peptide, a large number of cells died (Figure 1A), and the cell activity was better than the ADSCs in the concentration intervention of the bee toxin peptide concentration intervention in the group B and group C, but it Cell activity still shows gradient decreased gradient and lower than the cell activity of the control group. When ADSCs intervenes under the concentration intervention of the bee toxin peptide of the group D, it can be found that its overall cell activity is the same as the control group cell activity. The cell activity is better. Obviously differences, and found that when the concentration of bee toxin peptide was 0.04 mg/L and 0.08 mg/L, its cell activity was on the rise, which were 1.04 times and 1.08 times the cell activity of the control group, and the concentration of bee toxin peptide was 0.32 at 0.32 At MG/L, its cell activity decreases compared with the cell activity of the control group (Figure 1B-1D). When ADSCs intervenes in the concentration intervention of the group D bee toxin, its ADSCs has a good overall state, the cells have no obvious death, the cell activity is good, and when the concentration of bee toxin peptide is 0.04 mg/L and 0.08mg/L, its cell activity is available Rising trend is better than the cell activity of the control group.

**Figure 1:**
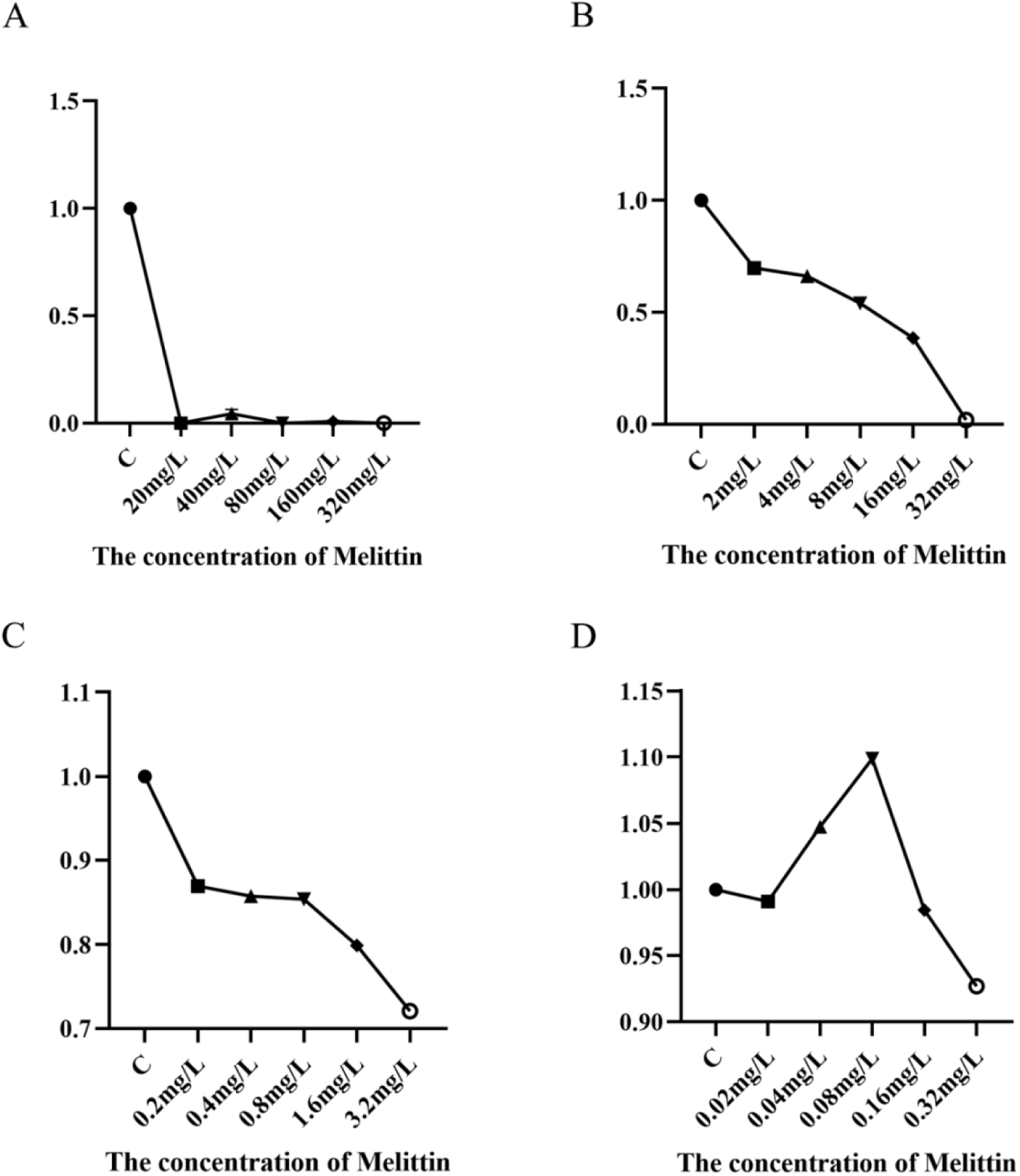
Melittin in different concentration groups interferes with ADSCs for 48 hours. (A) The effect of Melittin concentration range (20 mg/L, 40 mg/L, 80 mg/L, 160 mg/L, 320 mg/L) in group A on the cell activity of ADSCs. (B) The effect of Melittin concentration range (2 mg/L, 4 mg/L, 8 mg/L, 16 mg/L, 32 mg/L) in Group B on the cell activity of ADSCs. (C) The effect of Melittin concentration range in group C (0.2 mg/L, 0.4 mg/L, 0.8 mg/L, 1.6 mg/L, 3.2mg/L) on ADSCs cell activity, (D) Melittin concentration range in group D (0.02 mg/L, 0.04 mg/L, 0.08 mg/L, 0.16 mg/L, 0.32mg/L) on the cell activity of ADSCs. P<0.05.

### 3.2 Appropriate concentration of bee toxin peptide can enhance the expression of muscle differentiation in ADSCs

In order to explore the expression related to the deeper differentiation of ADSCs after the bee toxin intervention intervention, the control group and 0.04mg/L, 0.08mg/L, 0.16mg/L were selected. The protein expression level (Figure 2A-2C) is detected to be tested into MYOD, MYF5, MYH4. The experimental results show that the level of muscle differentiation related factors of ADSCs after melittin intervention is better than the level of protein expression in the control group. When the concentration of bee toxin peptide is 0.08 mg/L and 0.16mg/L, the correlation related factors of the muscle differentiation The expression level is better than the remaining two groups, and when the concentration of melittin is 0.08 mg/L, the expression level of ADSCs’s muscle differentiation related factors is best expression. Times, 1.24 times and 1.67 times, and when the concentration of bee toxin peptide is 0.04 mg/L, the level of ADSCs’s muscle differentiation related factors is time to 0.08mg/L. 1.27 times, 1.08 times and 1.24 times.

**Figure 2:**
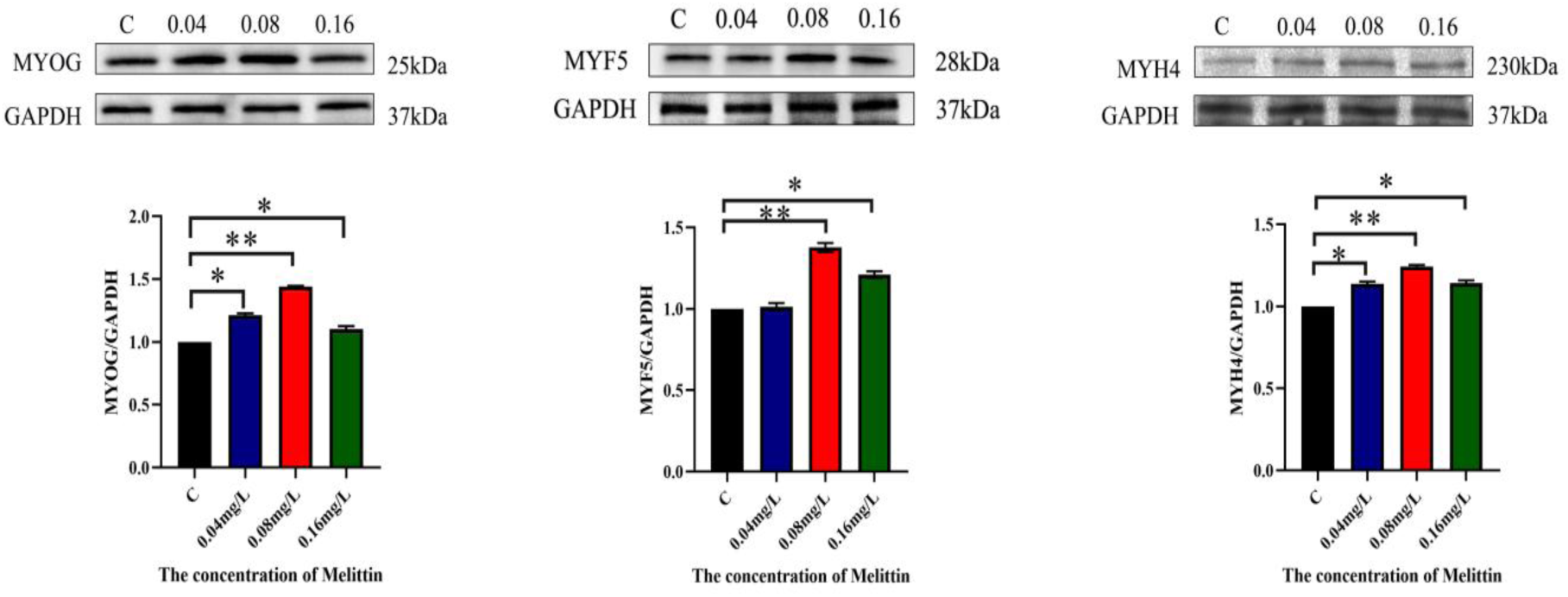
The effect of concentration of concentration melittin on cell regeneration factor. (A): MYOD, MYF5, MYH4 detection and protein marks analysis of protein levels. (B): MYOG, MYF5, MYH4 quantitative analysis of protein levels. *P <0.05, ** P <0.01, *** P <0.001, **** P <0.0001.

### 3.3 Bee toxin peptide can improve the expression of muscle differentiation in ADSCs

In order to explore the effects of melittin on atrophy cells, the cell atrophic model is established through the seamisonon atrophy, and the control group (CONTROL, C), the Melittin (M), and the dexamethasone, D), Dexamethasone+Melittin (D+M), interfere with 12h in the ground Cymet (10umol/L), add melittin intervention for 48h, and use Western-blot to detect it to Somathered related factors MYOG, MYOD,MYF5, MYH4 (Figure 3A-3B). The group is 1.10 times, 1.36 times, 1.34 times, and 1.24 times. The experimental results show that melittin can increase the related factors of the synthrodia into the synthydividation related factors.

**Figure 3:**
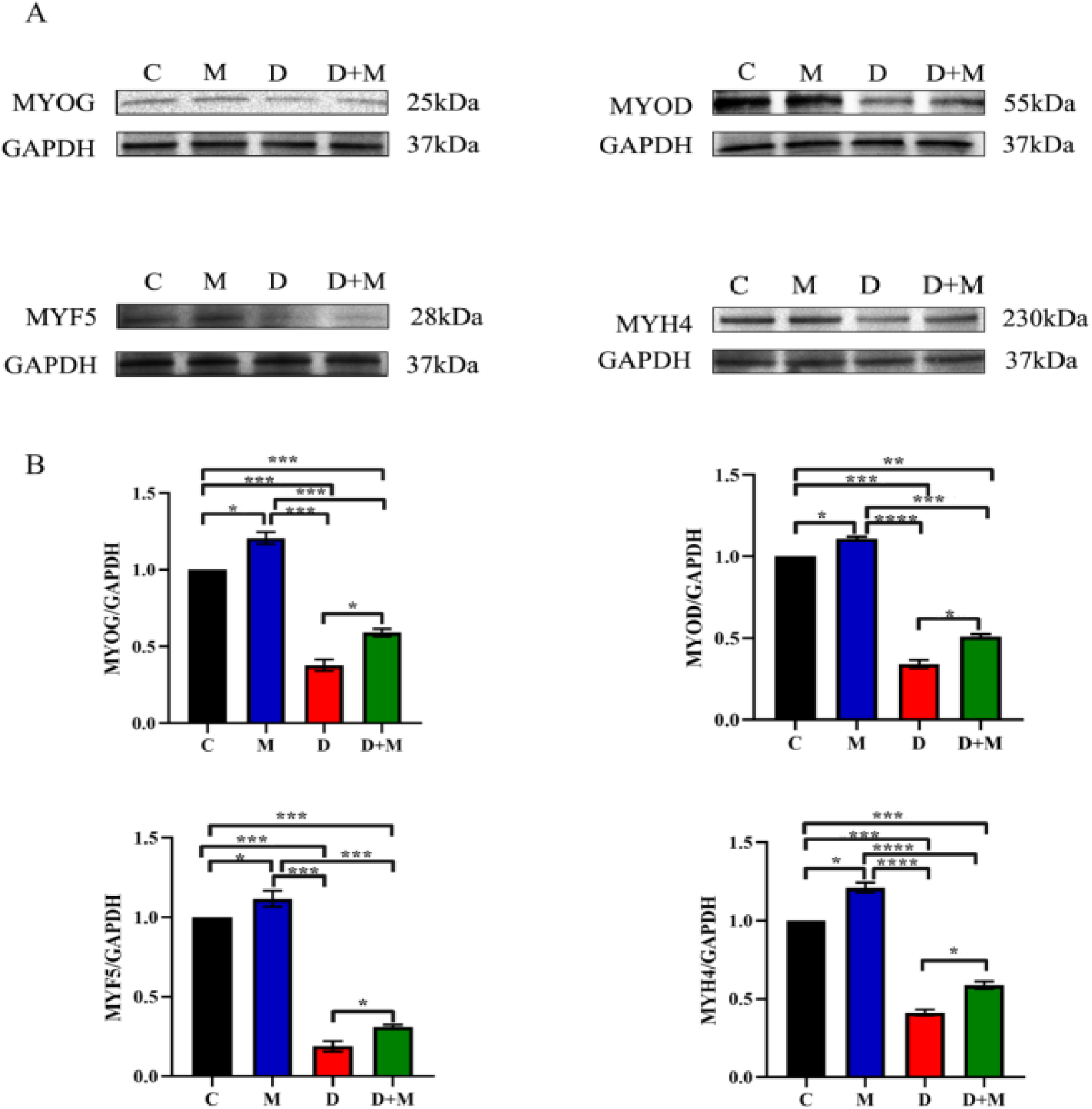
The effects of bee toxin peptide on the correlation factors of the innerized division of atrophy cells. Dicemethon induced ADSCs atrophy 12h and gives melittin intervention 48h. (A): MYOG, MYF5, MYOD, MYH4 detection and protein marks analysis. (B): MYOG, MYF5, MYOD, MYH4 quantitative analysis. *P <0.05, ** P <0.01, *** P <0.001, **** P <0.0001.

In order to observe the effects of bee toxin peptide on atrophy cells, the cells are also divided into four groups, C groups, groups, group D, D+M group, and detect MYOD, MYOG (Figure 4A-4B) in the immune fluorescent method. After the D+M group induces the atrophic cells, the expression of the correlation-related factors MYOD and MYOG is higher than the atrophic cells (4C-4D) that is higher than the D group. The results of the study show that melittin can improve the expression of muscle differentiation related factors in the atrophic ADSCs.

**Figure 4:**
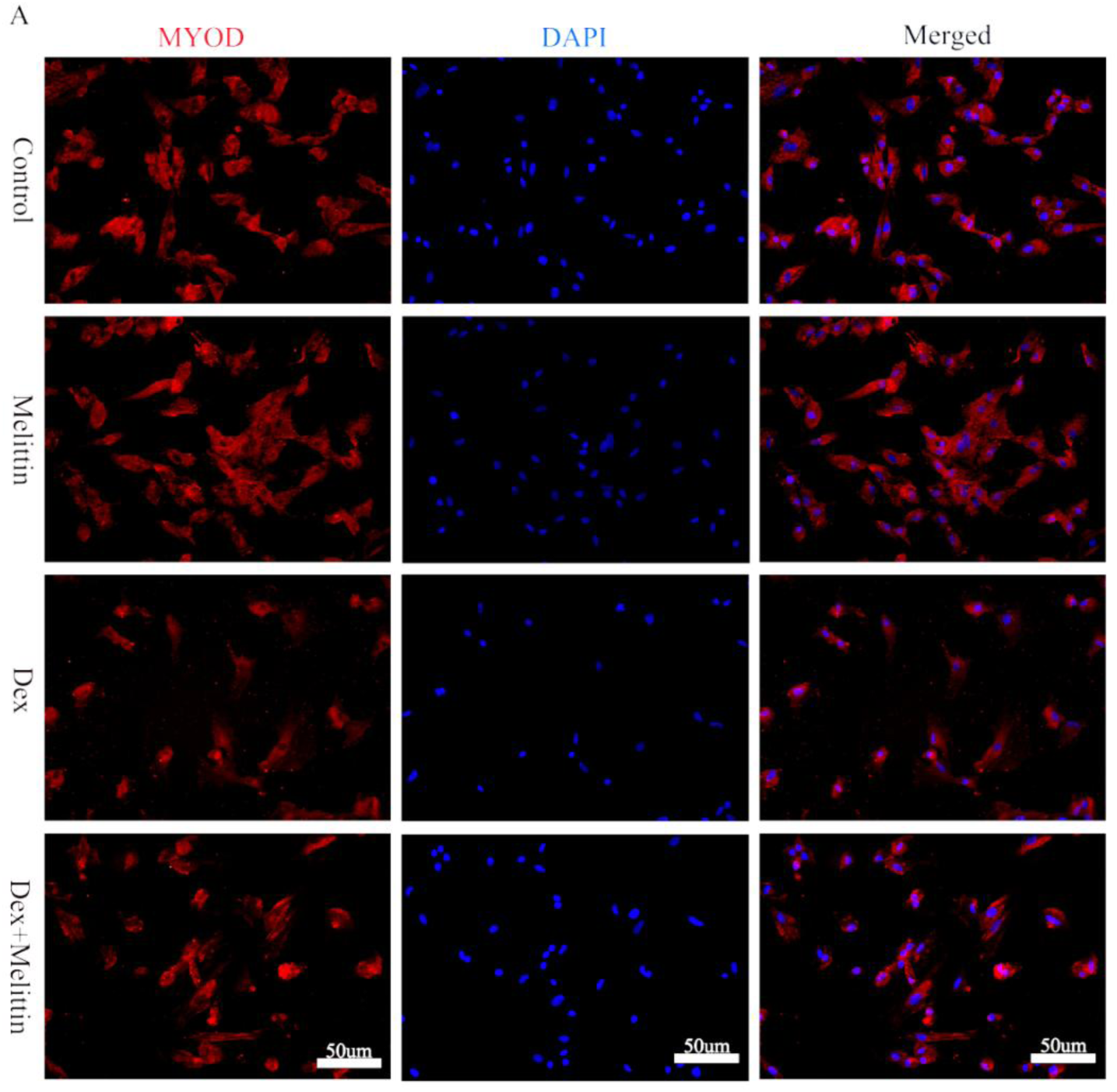
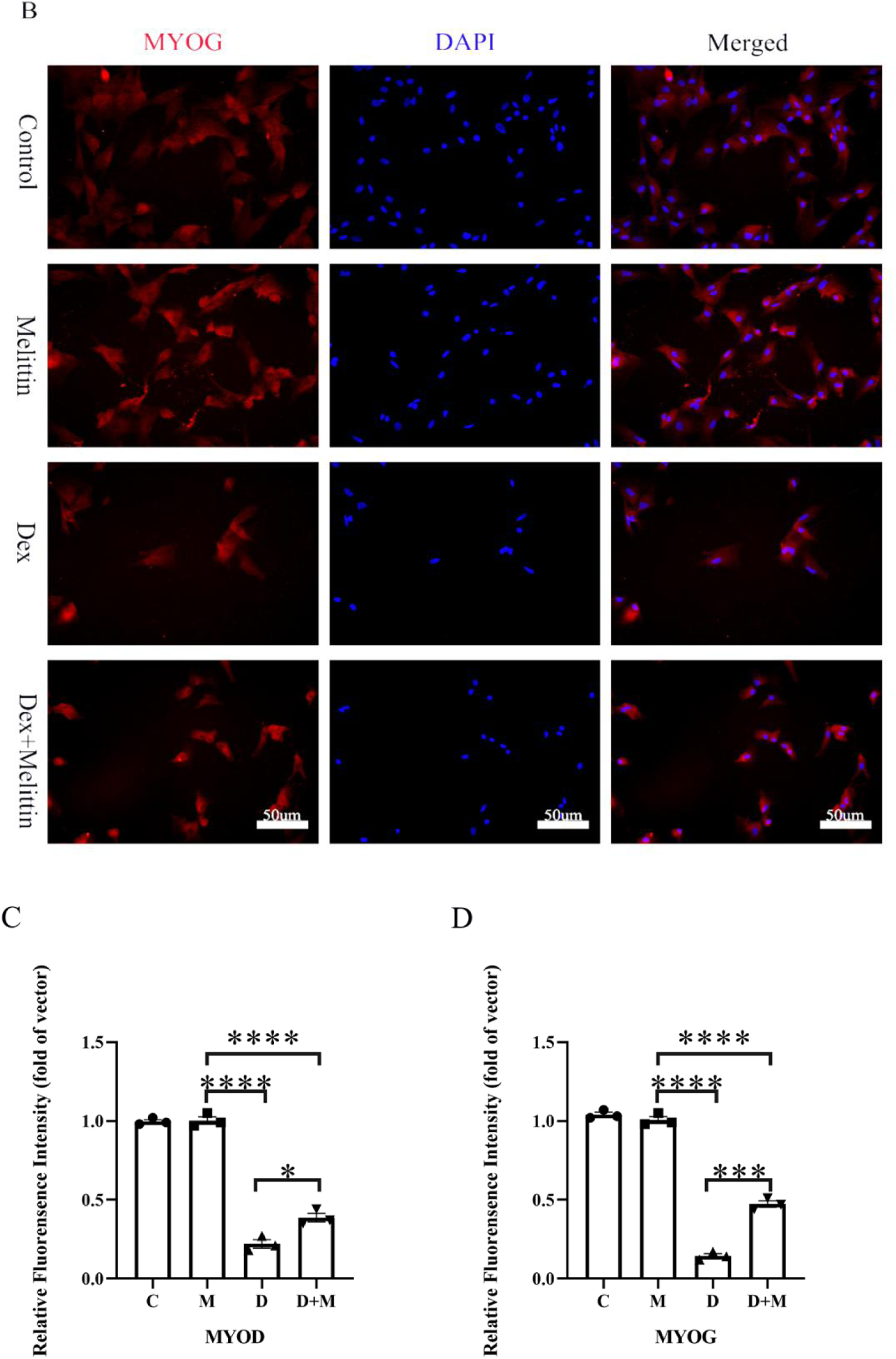
Immune fluorescence explores bee toxic induction of ADSCs formation. (A): The representative image and quantitative data of MYOG (red). (B): MYOD (red) represents the image. (C-D): Cells are relatively fluorescent. *P <0.05, ** P <0.01, *** P <0.001, **** P <0.0001.

### 3.4 Bee toxic peptide can promote ADSCs muscle differentiation through the p38-MAPK signaling pathway

According to the above results, melittin can significantly enhance the effect of ADSCs’s proliferation and muscle diversion, but its molecular mechanism is unclear. Through consulting relevant literature, p38-MAPK participates in a series of muscle differentiation biological processes. Therefore, the level of phosphorylation of p38-MAPK is the effect of beexing peptide to promote ADSCs.

In order to explore the effects of bee toxin peptide on the p38 phosphate in the process of ADSCs’s muscle division, the cell atrophy model is established through the seamisonon induction of ADSCs atrophy, and the control group (CONTROL, C), Melittin (M), DEXAMETHASONE (D), intervene in the ADSCs Gloson (10umol/L) for 12h, add melittin intervention for 48h, and use Western Blot to detect p38 and p-p38 protein (Figure 5A-5C). The results show that the expression level of the p-p38 expression of the D+M group is higher than the expression level of the p-p38 group p-p38, indicating that the activity form of the p-p38 is enhanced and promoted differentiation. Improve the activity of P-P38, and the level of p-p38 in group M is higher than that of group C. The results showed that melittin activated the P38-MAPK signaling pathway and promoted the decentralization of ADSCs.

**Figure 5:**
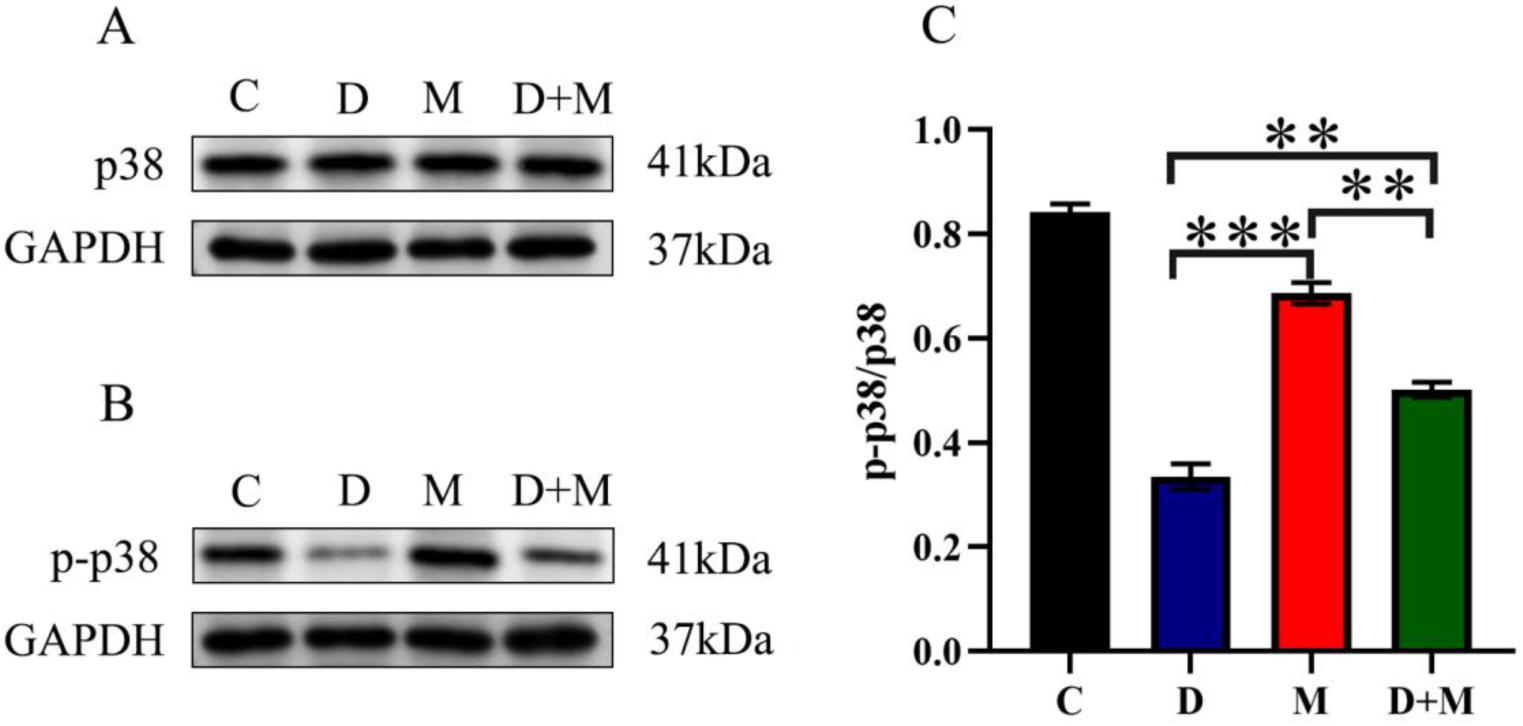
The effects of bee toxin peptide on the p38 phosphorylation during the splitting of ADSCs. Dicemethon induced ADSCs atrophy 12h and gives melittin intervention 48h. (A): p38, p-p38, GAPDH detection and protein marks analysis of protein levels. (B): quantitative analysis of p38, p-p38, and Gapdh of protein levels. *P <0.05, ** P <0.01, *** P <0.001, **** P <0.000

### 3.5 ADSCs intervention across Melittin can enhance the restoration of skeletal muscle morphology of rats

In order to explore the effects of ADSCs after melittin intervention on skeletal muscle atrophy, 48 SD male rats (240 ±10g) were selected, randomly divided into (Control, C), damage group (Tears, T), stem cell therapy group (STEM Cells, SC), melittin combined with ADSCs therapy group (STEM Cells+Melittin, SC+M). Group C does not do any treatment, group T, SC group, and SC+M set up a mice skeletal muscle atrophy model, all from the left side of the rats on the left gang tendon, and mark it with surgical sutures. On the first day of the operation, The SC group of the muscle injection of ADSCs ((1 ×10^6^), and the ADSCs after the intervention of the melittin intervention from the SC+M group, repeat the above treatment every 7 days. Large Gangto tendon, extract muscle (Figure 6A-6B).

**Figure 6:**
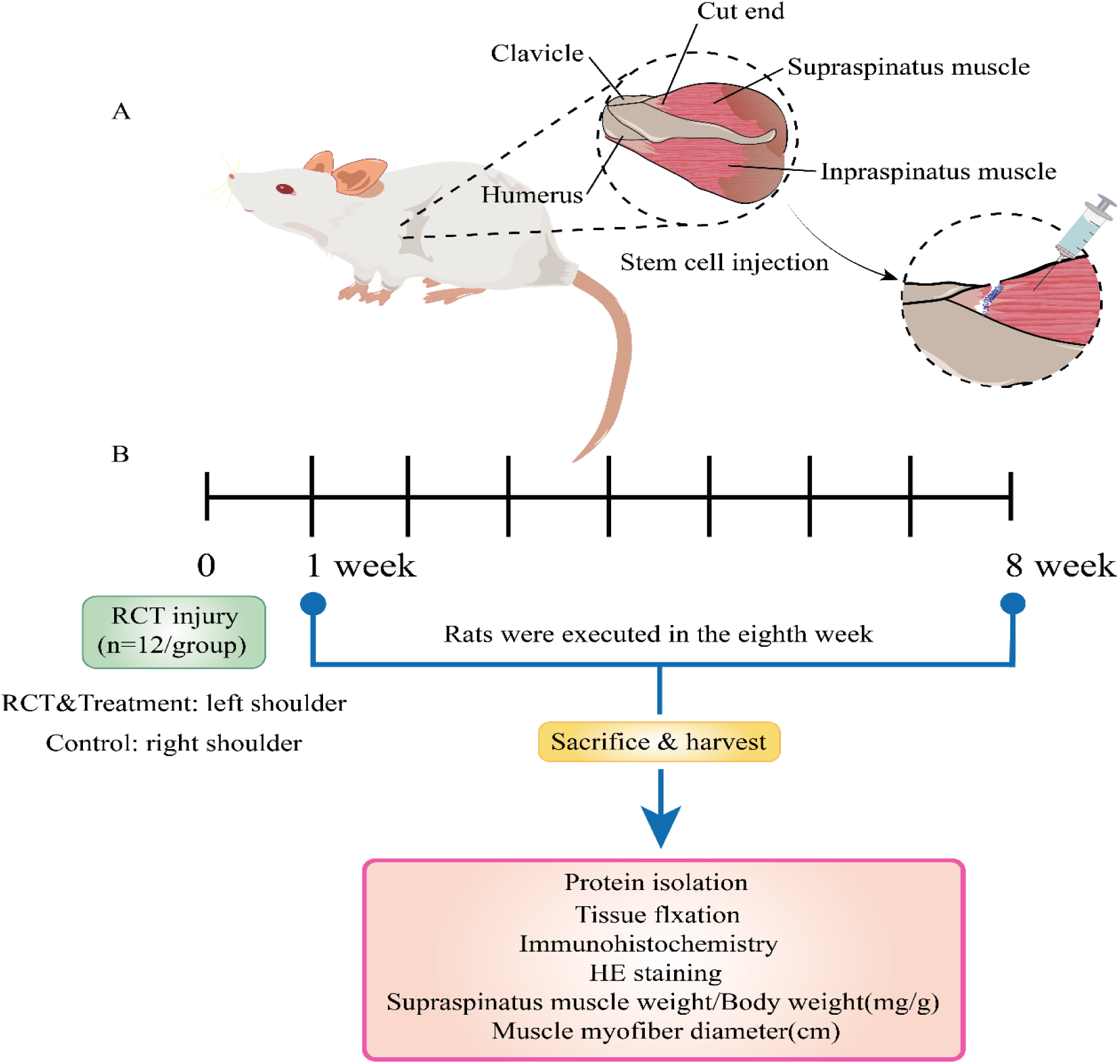
48 male SD mice are randomly divided into four groups: control group (C), damage group (T), stem cell therapy group (SC), beex peptide combined with stem cell therapy group (SC+M). (A): The establishment of the skeletal muscle atrophy of the rat. After the skin is cut, cut it vertically on the bougainvillea, exposing the shoulder sleeve tendon at the shoulder joint. The tendon cutting group completely cut off the upper of the Gang and the lower ligaine tendon, and sutured the sewing of the end. On the first day after surgery, ADSCs (about 1 ×10^6^) was injected with SC group, and ADSCs (approximately 1 ×10^6^) was injected with SC+M group SD rats in the SC+M group, and then repeated every seven days. The above treatment. (B): All rats are executed after eight weeks, separated the left gandan tendon, extract muscles, and performs protein extraction, tissue slices, immunohistochemistry, HE dyeing and other related testing.

In order to study the impact of ADSCs after melittin intervention on the morphological changes of muscle tissue, H & E dyed and immunohistochemicals were used for related testing (Figure 7A-7F). Compared with SC group skeletal muscle atrophy rats, the average cross-sectional area (Cross-SETIONALA MEAN) of SC+M group is higher than the SC group, and the muscle fiber form is not significantly different from group C. The intervention ADSCs is better than simply stem cell therapy in restoration of muscle fiber structure, while inhibiting skeletal muscle atrophy and improving the muscle tissue after injury. The result of immunohistochemicals shows that the MYOD and MYOG protein expression of MYOD and MYOG protein compared with the T group and SC+M groups. At the same time, the weight of the tendon of the upper of Rat Orthoga and the diameter of muscle fiber were measured. All found that the muscle weight and diameter of the SC+M group were better than the SC mouse (Figure 7G-7H). The experimental results show that ADSCs intervention intervention with melittin can enhance the recovery of skeletal muscle morphology of rats.

**Figure 7:**
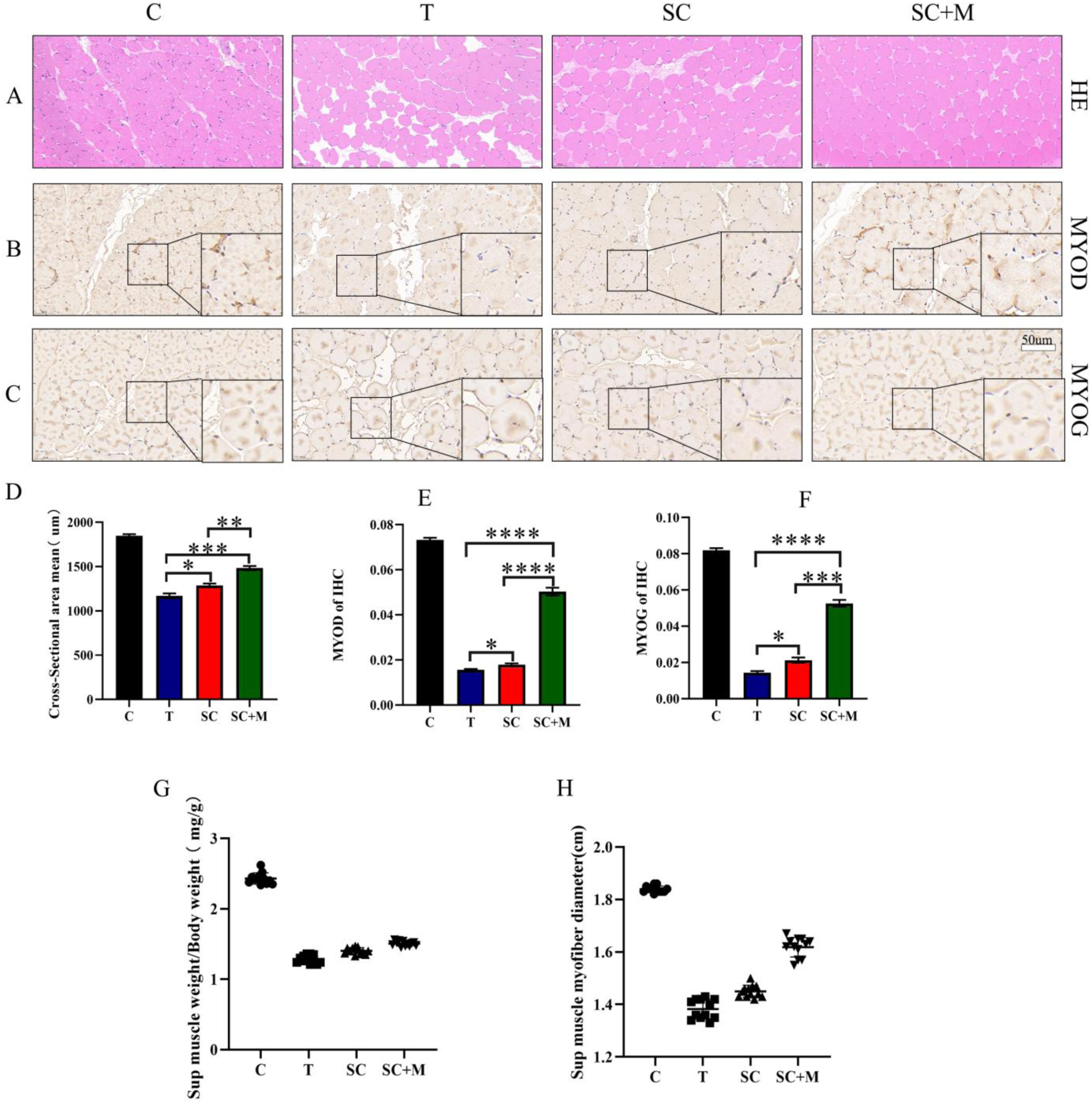
(A) Su Mu Jing and Yi Hong dye the slices of tendon in Rat Gang. (B) MYOD immunine dyeing slices. (C) MYOG immunine dyeing section. (D) HE dyeing rats Okama’s upper muscle cross section average (total area of muscle fiber cross section/number of muscle fibers). (E-F): Optical results of immunozoic chromators. (G): Detecting muscle weight, the weight of the upper muscle weight to SD rats. (H): Group C, Group T, Group SC,Group SC+M Gang upper muscle fiber diameter. *P <0.05, ** P <0.01, *** P <0.001, **** P <0.0001.

### 3.6 ADSCs after Melittin intervention can enhance the treatment of rats’ skeletal muscle atrophy, and strengthen the expression of related factors related to muscle differentiation

In order to explore the expression of muscle differentiation in muscle tissue, after extracting muscle protein, Western-blot is detected into MYOG, MYF5, MYOD, MYH4. The experimental results show that the expression of SC group and SC+M composition of muscle differentiation related factors is better than the T group. The SC+M group MYOG and MYF5 protein expression are 1.51 times and 1.49 times of SC groups, and the MYOD, and the SC+M group’s MYOD, and the MYOD in groups, and the MYOD, and the MYOD of the SC+M group, and the MYOD in the SC+M group, and the MYOD in the SC+M group. The expression of MYH4 is 1.42 and 1.47 times of the SC group (Figure 8A-8B). It shows that ADSCs intervention after melittin can enhance the treatment of rats’ skeletal muscle atrophy, and strengthen the expression of related factors of muscle differentiation.

**Figure 8:**
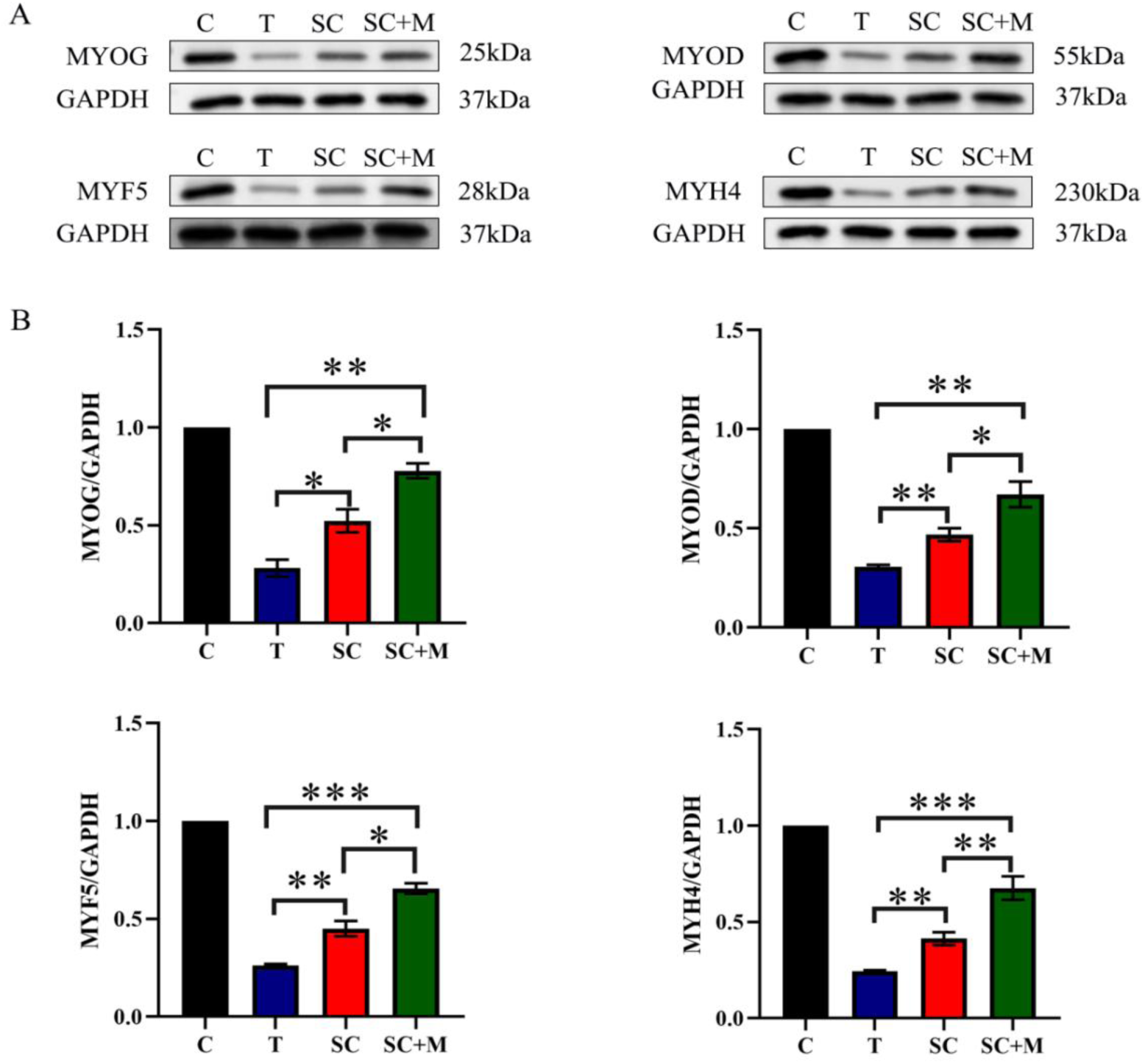
Western-blot is detected into muscle differentiation related factors. (A) MYOG, MYOD, MYF5, MYH4 of protein levels. (B) MYOG, MYOD, MYF5, MYH4 quantitative analysis of protein levels. *P <0.05, ** P <0.01, *** P <0.001, **** P <0.0001.

## Disscussion

Skeletal muscle atrophy is often caused by failure to recover in time after injury. When the damage is severe enough to affect muscle function, physical exercise will be limited. In recent years, people have made considerable efforts to continuously explore and develop inhibitory of skeleton muscle atrophy drugs and treatment methods. This study has confirmed that melittin can use ADSCs’s innate regeneration potential to promote it into muscle division, and at the same time reveal the role of melittin the treatment of skeletal muscle atrophy under pathological conditions.

A large number of studies have shown that ADSCs has excellent potential in the treatment of skeletal muscle atrophy, but the simple ADSCs treatment effect is not good. Past studies have shown that only about 15%of ADSCs can be completely differentiated to in the differentiation medium ^[7]^, and if it is directly, it will be directly. ADSCs transplants to target organs will have defects such as ADSCs low survival rate and low partialization efficiency ^[18]^. Therefore, pre-processing of stem cell transplantation can be pre-mounted to improve stem cell survival rate and differentiation efficiency into a potential treatment strategy.

MYOG, MYF5, and MYOD are members of MYOGenic regulatory factor family (MRF). MRF is important in guiding muscle satellite cell function and preventing skeletal muscle atrophy^[19]^. At the same time, MYF5 and MYOD are considered to be the determined factors of MYOGenicity. The ability to induce fibroblasts and various non-skeletal muscle cells into muscle differentiation, while re-programming the ability of other cells as skeletal muscle cells, and because of its expression in early muscle spectrum, MYOD is usually called the main regulation. Factors ^[22-23]^, and MYF5 is first expressed during the development of embryo muscle, and the transformation of muscle stem cells with muscle satellite cells with muscle cells is activated ^[24]^. And it is necessary for the ending of skeletal muscle cells in the embryo ^[25]^.

In cell experiments, the ADSCs after melittin intervention, the expression related factor MYOG, MYF5, and MYOD in MYOGenization is significantly higher than the SC group, and improved the expression of the skin differentiation related factors of the shrinking ADSCs. The result of immunofluorescence test also shows that MYOD and MYOG signals in the D+M group are higher than that of group D cells. It shows that melittin can improve the expression of muscle differentiation by improving ADSCs muscle differentiation efficiency, enhance the expression of related factors of muscle differentiation, and then inhibit muscle atrophy.

At the same time, animal experiments have shown that compared with the SC group, ADSCs after melittin intervention treats skeletal muscle atrophy rats, and its muscle differentiation related factors are significantly better than the SC group. Skeletal muscle atrophy of skeletal muscle atrophy of stem cells after peptide intervention is significantly improved in muscle weight, muscle fiber diameter, and muscle fiber cross-sectional area than SC rats. At the same time It is significantly better than the SC group, so there may be a new fiber formation in the treatment process to inhibit skeletal muscle atrophy ^[9]^.

In this study, compared with the SC group, the skeletal muscle atrophy of skeletal muscle atrophy of skeletal treatment after melittin intervention has a significant increase in muscle differentiation related factors, including the essential factor MYOG and maintaining muscle quality for muscle differentiation The number of fibrous quantity MYH4 ^[26]^, as well as MYOD that can convert part of fibroblasts and other types of cells into skeletal muscle^[27-28]^. MYOD, MYH4, and MYF5 have similar functions and overlap in terms of guidance into muscle differentiation, which plays an important role in promoting ADSCs MYOGenous differentiation and improving ADSCs activity ^[21]^.

The results of the above experiments show that bee toxin peptide can enhance the value-added and muscle division of ADSCs, but the molecular mechanism is unclear. It can be seen by consulting the relevant information. Growth and repair, and the original muscle transcription of MYOD class only has the p38-Mapk pathway directly affects it ^[29]^. At the same time, in the process of formation, skeletal muscle cell proliferation can be regulated with the p38-Mapk signal to be adjusted and fused to form a muscle tube ^[30]^. Therefore, speculation is speculated that the bee toxin peptide has participated in the p38-Mapk signaling pathway to promote ADSCs muscle differentiation. Therefore, the cell atrophy model is established, the control group (C), the bee toxic peptide group (M), the dexamethasone group (D), the dexamethasone+beex peptide group (D+M), and the ADSCs places the ground, After 10umol/L) intervention 12h, add melittin intervention for 48h, and use Western-BLOT to detect p38 and p-p38 protein. The experimental results show that melittin can enhance the P-38 phosphate process, thereby improving muscle division.

Skeletal muscle atrophy is affected by multiple factors. At present, there is no exact therapy to inhibit skeletal muscle atrophy. At the same time, a single drug intervention cannot achieve the expected treatment effect. Therefore Stem cells treat the muscle atrophy after bone injury of rats, trying to provide experiments and theoretical basis for the therapy. Because of its own quantity advantages, ADSCs is easy to obtain, and its inherent good differentiation potential and self-update ability have shown excellent application potential in the direction of skeletal muscle atrophy ^[26]^. However, in the process of applying ADSCs to treat skeletal muscle atrophy, the main points that can effectively exert their therapeutic effects are the effective proliferation of ADSCs and the efficiency of muscle division.

Drug safety is a major problem in clinical application of beex peptide, which may cause drug allergic reactions ^[27]^, and melittin is considered to be one of the bee toxic components that cause allergic reactions. However, research shows that bee toxin peptides can be used safely. In the research of toxicology, the LD 50 of bee toxic peptide to SD rats is greater than 30.0 mg/kg ^[28]^. In another 13-week toxicology research, Repeat muscle injection bee toxin peptide until 0.28 mg/kg (far higher than clinical dosage), no significant toxicity ^[31]^. In the current research, we have not observed any melittin in animal experiments. The effect of toxicology, at the same time, melittin has been widely used in anti-inflammatory, in the clinical treatment of cancer ^[32]^, comprehensive consideration, the clinical use of melittin is relatively safe, and the purified beex peptide is easy to control in actual use in actual use. Advantages of the treatment dose.

There are still relevant limitations in this study. First of all, when the drug is injected into the damage to the muscles of the rats, it is difficult to accurately locate stem cells to confirm whether its stem cells have reached damaged skeletal muscle tissue. At the same time Precise marks. Secondly, we cannot fully explore the effect of stem cells after melittinintervention. At the same time, the sample amount of the experimental animal is relatively low. Therefore, it is necessary to continuously optimize the above problems and in-depth research.

## Conclusion

The results of this study indicate that melittin can promote the MYOGenic differentiation of ADSCs and inhibit cell atrophy of ADSCs. Stem cells treated with bee venom peptide intervention can improve the recovery of muscle mass, muscle fiber diameter, and muscle fiber cross-sectional area in rats with skeletal muscle atrophy compared to rats treated with stem cells alone. Secondly, melittin can activate the p38-MAPK signaling pathway and promote the MYOGenic differentiation of ADSCs. Therefore, the above results provide insights into MYOGenic differentiation and a potential alternative strategy for treating skeletal muscle atrophy.

## Ethics Declaration

The institutional Ethics Committee of SuBei People’s Hospital approved this study and included experimental procedures. All animal housing and experiments were conducted strictly with the institution’s rules for the care and use of laboratory animals.

## Consent to publication

All authors approved the final manuscript and the submission to this journal.

## Authors Contributions

Fei W.Y and Li M.J. designed the studies, performed the procedures, analyzed the results, wrote the paper, wrote the editing review, dealt with methodology and validation, performed the data analysis, and conducted the study as a leader; conceptualization, Fei W.Y; Li M.J.; Yang Y.X; Zhang, J; methodology, Meng X.J; Liu W.K; Wang S.G; Liu D.W;Software, Dang M.B; Yang, J; Dai X.M;validation, Li M.J.; Yang Y.X; Zhang, J; formal analysis, Yang, J; Dai X.M; Bao, T; Wang, Y; investigation, Bao, T, and Wang, Y; resources, Fei W.Y and Li M.J.; data curation, Fei W.Y, and Li M.J.; writing—original draft preparation, Fei W.Y, and Li M.J.; writing—review and editing, Yang Y.X and Zhang, J; visualization, Fei W.Y; supervision, Fei W.Y; Li M.J.; Yang Y.X;project administration, Fei W.Y;funding acquisition, Fei W.Y; Li M.J. All authors have read and agreed to the published version of the manuscript.

## Funding

The funding for this research institute is provided by—

1. Load porous Se@SiO2 Experimental Research on Nanomaterials and BMP-2 Relief Hydrogel Combined with Adipose Stem Cells to Promote Shoulder Sleeve Tendon bone Healing, National Orthopaedic and Sports Rehabilitation Clinical Medical Research Center, No. 2021-NCRC-CXJJ-PY-07;
2. Experimental study on the application of PLGA/hydroxyapatite 3D printing scaffold combined with BMP-2 in the repair of rotator cuff tendon-bone interface damage, Jiangsu Provincial Health Commission project, No. M2021042;
3. Experimental study on the evaluation of stem cell therapy for degenerative rotator cuff injury using superparamagnetic iron oxide tracing technology, Yangzhou Social Development Project, No. YZ2021084.
4. The Science and Technology Bureau of Yangzhou City(Project number: YZ2021084).

## Available data and materials

The article’s data and online supplementary material are available in the paper.

## Acknowledgments

We appreciate the contribution of Dr. Fei Wenyong, the author, to this experiment.

## References

1. Liang JL, Xie JF, Wang CY, Chen N (2020) [Regulatory roles of microRNAs in sarcopenia and exercise intervention]. Sheng Li Xue Bao 72:667–676

2. Cohen S, Nathan JA, Goldberg AL (2015) Muscle wasting in disease: molecular mechanisms and promising therapies. Nat Rev Drug Discov 14:58–74

3. Wang W, Shen D, Zhang L, Ji Y, Xu L, Chen Z, Shen Y, Gong L, Zhang Q, Shen M, Gu X, Sun H (2021) SKP-SC-EVs Mitigate Denervated Muscle Atrophy by Inhibiting Oxidative Stress and Inflammation and Improving Microcirculation. Antioxidants (Basel) 11:66

4. Talarek JR, Piacentini AN, Konja AC, Wada S, Swanson JB, Nussenzweig SC, Dines JS, Rodeo SA, Mendias CL (2020) The MRL/MpJ Mouse Strain Is Not Protected From Muscle Atrophy and Weakness After Rotator Cuff Tear. J Orthop Res 38:811–822

5. Simon CB, Coronado RA, Greenfield WH 3rd, Valencia C, Wright TW, Moser MW, Farmer KW, George SZ (2016) Predicting Pain and Disability After Shoulder Arthroscopy: Rotator Cuff Tear Severity and Concomitant Arthroscopic Procedures. Clin J Pain 32:404–410

6. Chen K, Yin S, Wang X, Lin Q, Duan H, Zhang Z, Chang Y, Gu Y, Wu M, Wu N, Liu C (2020) Effect of extracorporeal shock wave therapy for rotator cuff tendonitis: A protocol for systematic review and meta-analysis. Medicine (Baltimore) 99:e22661

7. Tornero-Esteban P, Hoyas JA, Villafuertes E, Rodríguez-Bobada C, López-Gordillo Y, Rojo FJ, Guinea GV, Paleczny A, Lópiz-Morales Y, Rodriguez-Rodriguez L, Marco F, Fernández-Gutiérrez B (2015) Efficacy of supraspinatus tendon repair using mesenchymal stem cells along with a collagen I scaffold. J Orthop Surg Res 10:124

8. Moussa MH, Hamam GG, Abd Elaziz AE, Rahoma MA, Abd El Samad AA, El-Waseef D, Hegazy MA (2020) Comparative Study on Bone Marrow-Versus Adipose-Derived Stem Cells on Regeneration and Re-Innervation of Skeletal Muscle Injury in Wistar Rats. Tissue Eng Regen Med 17:887–900

9. Jiang R, Wang M, Shi L, Zhou J, Ma R, Feng K, Chen X, Xu X, Li X, Li T, Sun L (2019) Panax ginseng Total Protein Facilitates Recovery from Dexamethasone-Induced Muscle Atrophy through the Activation of Glucose Consumption in C2C12 Myotubes. Biomed Res Int 2019:3719643

10. Liu L, Hu R, You H, Li J, Liu Y, Li Q, Wu X, Huang J, Cai X, Wang M, Wei L (2021) Formononetin ameliorates muscle atrophy by regulating myostatin-mediated PI3K/Akt/FoxO3a pathway and satellite cell function in chronic kidney disease. J Cell Mol Med 25:1493–1506

11. Zhang J, Zheng J, Chen H, Li X, Ye C, Zhang F, Zhang Z, Yao Q, Guo Y (2022) Curcumin Targeting NF-κB/Ubiquitin-Proteasome-System Axis Ameliorates Muscle Atrophy in Triple-Negative Breast Cancer Cachexia Mice. Mediators Inflamm 2022:2567150

12. Shin MJ, Shim IK, Kim DM, Choi JH, Lee YN, Jeon IH, Kim H, Park D, Kholinne E, Yang HS, Koh KH (2020) Engineered Cell Sheets for the Effective Delivery of Adipose-Derived Stem Cells for Tendon-to-Bone Healing. Am J Sports Med 48:3347–3358

13. Kim S, Kim K, Park J, Jun W (2021) Curcuma longa L. Water Extract Improves Dexamethasone-Induced Sarcopenia by Modulating the Muscle-Related Gene and Oxidative Stress in Mice. Antioxidants (Basel) 10:1000

14. Maulet Y, Brodbeck U, Fulpius BW (1982) Purification from bee venom of melittin devoid of phospholipase A2 contamination. Anal Biochem 127:61–67

15. Zhang Y, Yu DD, Cui DH, Liao X, Guo H (2018) [Report quality evaluation of systematic review or Meta-analysis published in China Journal of Chinese Materia Medica]. Zhongguo Zhong Yao Za Zhi 43:1254–1260

16. Lee JE, Shah VK, Lee EJ, Oh MS, Choi JJ (2019) Melittin - A bee venom component - Enhances muscle regeneration factors expression in a mouse model of skeletal muscle contusion. J Pharmacol Sci 140:26–32

17. Zammit PS (2017) Function of the myogenic regulatory factors Myf5, MyoD, Myogenin and MRF4 in skeletal muscle, satellite cells and regenerative myogenesis. Semin Cell Dev Biol 72:19–32

18. McClure MJ, Garg K, Simpson DG, Ryan JJ, Sell SA, Bowlin GL, Ericksen JJ (2016) The influence of platelet-rich plasma on myogenic differentiation. J Tissue Eng Regen Med 10:E239–249

19. Ceafalan LC, Popescu BO, Hinescu ME (2014) Cellular players in skeletal muscle regeneration. Biomed Res Int 2014:957014

20. Weintraub H, Davis R, Tapscott S, Thayer M, Krause M, Benezra R, Blackwell TK, Turner D, Rupp R, Hollenberg S, et a (1991) The myoD gene family: nodal point during specification of the muscle cell lineage. Science 251:761–766

21. Comai G, Tajbakhsh S (2014) Molecular and cellular regulation of skeletal myogenesis. Curr Top Dev Biol 110:1–73

22. Choi J, Costa ML, Mermelstein CS, Chagas C, Holtzer S, Holtzer H (1990) MyoD converts primary dermal fibroblasts, chondroblasts, smooth muscle, and retinal pigmented epithelial cells into striated mononucleated myoblasts and multinucleated myotubes. Proc Natl Acad Sci U S A 87:7988–7992

23. Chan SS, Kyba M (2013) What is a Master Regulator. J Stem Cell Res Ther 3:114 [pii]

24. Hasty P, Bradley A, Morris JH, Edmondson DG, Venuti JM, Olson EN, Klein WH (1993) Muscle deficiency and neonatal death in mice with a targeted mutation in the myogenin gene. Nature 364:501–506

25. Megeney LA, Rudnicki MA (1995) Determination versus differentiation and the MyoD family of transcription factors. Biochem Cell Biol 73:723–732

26. Wright WE, Sassoon DA, Lin VK (1989) Myogenin, a factor regulating myogenesis, has a domain homologous to MyoD. Cell 56:607–617

27. Kim SY, Kim MH, Cho YJ (2018) Different clinical features of anaphylaxis according to cause and risk factors for severe reactions. Allergol Int 67:96–102

28. Zeng C, Shi H, Kirkpatrick LT, Ricome A, Park S, Scheffler JM, Hannon KM, Grant AL, Gerrard DE (2021) Driving an Oxidative Phenotype Protects Myh4 Null Mice From Myofiber Loss During Postnatal Growth. Front Physiol 12:785151

29. Qi R, Liu H, Wang Q, Wang J, Yang F, Long D, Huang J (2017) Expressions and Regulatory Effects of p38/ERK/JNK Mapks in the Adipogenic Trans-Differentiation of C2C12 Myoblasts. Cell Physiol Biochem 44:2467–2475

30. Korb A, Tohidast-Akrad M, Cetin E, Axmann R, Smolen J, Schett G (2006) Differential tissue expression and activation of p38 MAPK alpha, beta, gamma, and delta isoforms in rheumatoid arthritis. Arthritis Rheum 54:2745–2756

31. Kang H, Lim C, Kwon KR, Lee K (2014) Study of a 13-weeks, Repeated, Intramuscular Dose, Toxicity Test of Sweet Bee Venom in Sprague-Dawley Rats. J Pharmacopuncture 17:73–79

32. Sun M, Wu Y, Zhou Z, Liu S, Mao S, Li G (2023) Co-delivery of EGCG and melittin with self-assembled fluoro-nanoparticles for enhanced cancer therapy. Aging (Albany NY) 15:4875–4888

